# Rapid coordination of effective learning by the human hippocampus

**DOI:** 10.1101/2020.10.20.347831

**Authors:** James E. Kragel, Stephan Schuele, Stephen VanHaerents, Joshua M. Rosenow, Joel L. Voss

## Abstract

Although the human hippocampus is necessary for long-term memory, controversial findings suggest that hippocampal computations support short-term memory in the service of guiding effective behaviors during learning. We tested the counterintuitive theory that the hippocampus contributes to long-term memory through remarkably short-term processing, as reflected in the sequence of eye movements during encoding of naturalistic scenes. While viewing scenes for the first time, participants generated patterns of eye movements that reflected a shift from stimulus-driven to memory-driven viewing and signaled effective spatiotemporal memory formation. Hippocampal theta oscillations recorded from depth electrodes predicted this viewing pattern. Moreover, effective viewing patterns were preceded by shifts towards top-down influence of hippocampal theta on activity within cortical networks that support visual perception and visuospatial attention. The hippocampus thus supports short-term memory processing that coordinates perception, attention, and behavior in the service of effective spatiotemporal learning. These findings motivate re-interpretation of long-term memory disorders as reflecting loss of the organizing influence of hippocampal short-term memory on learning.

---

The hippocampus is essential for long-term memory (1) and memory-guided behaviors such as spatial navigation (2–4). For example, long-term memory guides visuospatial attention (5, 6), such that past experiences can influence the rapid (~2-5/s) saccadic eye movements needed to sample complex stimuli such as visual scenes (7–9). Yet, the role of the hippocampus in guiding visual sampling might be far more immediate, supporting online representations that emerge across sequential visual fixations and rapidly guide choices of where to look next (10, 11). Indeed, during the first exposure to complex stimuli, hippocampal lesions disrupt viewing patterns that reflect building a memory for the relations among distinct stimulus features (12, 13) and such viewing patterns correlate with hippocampal activity as measured via fMRI in healthy individuals (12, 14). Failure to effectively sample relations among distinct features via eye movements could also underlie impaired perception and short-term retention of complex stimuli identified following hippocampal lesions (15–18). These short-term behavioral deficits are surprising given the standard model of hippocampal involvement in long-term memory and suggest that long-term memory impairments could result from disrupted visual sampling during initial encoding. However, previous studies have been inconclusive in demonstrating that the hippocampus has a shortterm role in the rapid formation and online use of memory to guide viewing because those studies lacked the requisite spatial and temporal resolution.

We therefore used intracranial electroencephalography (iEEG) to provide temporally precise measurement of human hippocampal activity aligned to saccadic eye movements during scene memory formation (Fig. 1a). The goal was to test whether hippocampal activity recorded while participants (N=6) viewed novel scenes reflected the influence of the rapidly forming scene memory on effective eye-movement patterns that enhanced learning.

**Fig. 1.**
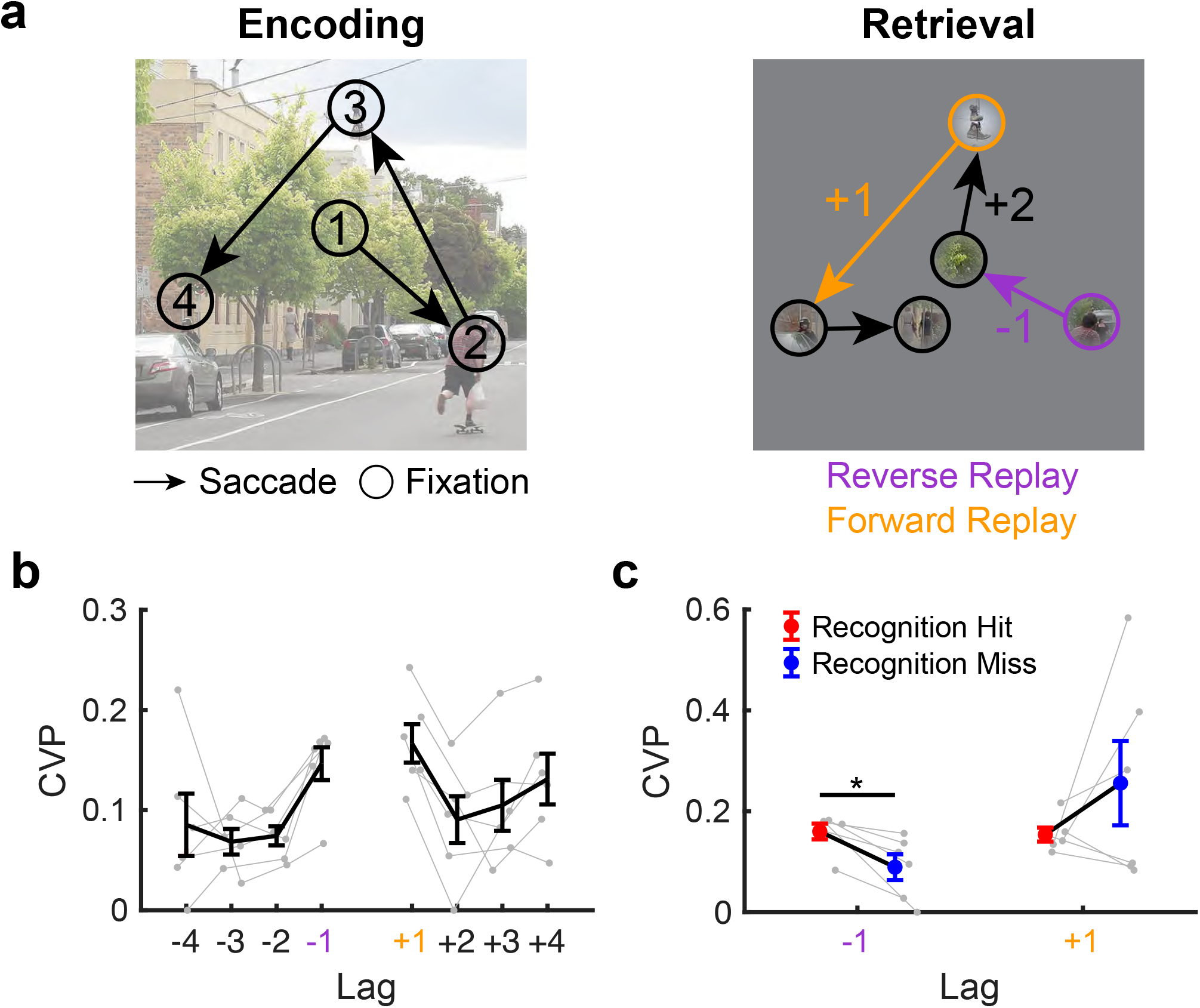
Fixation-sequence replay expresses spatiotemporal memory for scenes. **a**. During encoding, participants viewed scenes while eye-movement tracking recorded their visual fixation sequences. During retrieval, participants viewed repeated or novel scenes with gaze-contingent revealing of scene content, such that memory could guide viewing more so than peripheral perceptual information, which was masked. Retrieval fixations were coded based on their temporal distance (lag) in the encoding sequence. Spatiotemporal replay was identified in both forward (+1) and reverse (−1) directions. **b**. Participants replayed fixation sequences in forward and reverse directions, as indicated by the lag conditional viewing probability (lag-CVP) curve. **c**. Reverse replay was significantly greater on trials with accurate recognition responses. Error bars denote SEM. *, *P* < 0.05. Dots indicate individual participants.

Spatiotemporal memory is hippocampal dependent (19, 20) and can be observed for scenes as the tendency for subjects to replay during retrieval the scene-specific sequences of visual fixations that they made during encoding (21).Participants demonstrated robust fixation-sequence replay (lag-CVP, Fig. 1b). As expected, there was a tendency to replay temporally proximal as opposed to distal eye movements (i.e., a contiguity effect) (*F*(3,5) = 5.4, *p* = 0.01, *η*^2^ = 0.52 [95% CI, 0.40–0.95]), with prominent forward (+1 lags) and reverse (— 1 lags) replay distinguished from longer lags that were not indicative of replay. Visual sampling was matched in forward and reverse directions (*F*(1,5) = 2.0, *p* = 0.22, *η*^2^ = 0.28 [95% CI, 0.02–0.77]), with comparable contiguity effects in each direction (*F*(3,15) = 0.5, *p* = 0.71, *η*^2^ = 0.08 [95% CI, 0.02–0.63]). Furthermore, spatiotemporal replay of fixation sequences was related to the accuracy of overt scene recognition judgments. Participants reliably discriminated repeated from novel scenes (Supplementary Fig. 1a, with above-chance discrimination sensitivity (mean *d*’ = 1.7 ± 0.3 SEM), *t*(5) = 5.0, *p* = 0.004, *g* = 2.0 (95% CI, 0.6–3.5). Recognized scenes had higher levels of reverse replay than unrecognized scenes (*F*(1,5) = 18.9, *p* = 0.01, *η*^2^ = 0.76 [95% CI, 0.67–0.98]), but this difference was not observed for forward replay (*F*(1,5) = 1.1, *p* = 0.34, *η*^2^ = 0.18 [95% CI, 0.0007–0.85]). Thus, participants exhibited spatiotemporal memory for scenes as both forward (+1) and reverse (—1) replay of fixation sequences that was associated with overall scene recognition, which is consistent with previous findings in adults without epilepsy (21).

Patterns of encoding fixations that reflected spatiotemporal memory formation were defined based on their prediction of fixation-sequence replay during retrieval. We focused on a viewing pattern that has been associated with hippocampal function and/or memory formation in multiple previous studies in humans and rodents (22, 23). This “revisitation” viewing pattern occurs when subjects return to fixate the previously viewed location as opposed to moving on to a new location (Fig. 1a). Revisitation occurred for a minority of overall encoding fixations (mean = 39% ± 0.02 SEM) indicating that this viewing pattern countered the dominant tendency within scene viewing to sample novel content (24). As hypothesized, revisitation enhanced spatiotemporal memory (Fig. 1c). Fixations to revisited locations were about twice as likely to be replayed later during retrieval than were other non-revisited locations in both the forward direction (M=0.15, SEM=0.03 versus M=0.08, SEM=0.01, respectively, *t*(5) = 4.2, *p* = 0.008, *g* = 1.2 95% CI, 0.3–2.5) and in the reverse direction (M=0.12, SEM=0.02 versus M=0.05, SEM=0.01, respectively, *t*(5) = 4.2, *p* = 0.009, *g* = 1.2 95% CI, 0.3–2.3). Locations were just as likely to be revisited during encoding of later recognized (M=40%, SEM=0.01) as compared to forgotten (M=38%, SEM=0.04) scenes, *t*(5) = 0.55, *p* = 0.61, *g* = 0.2 (95% CI, −0.7–1.1). Thus, revisitation fixations reflected effective visual sampling that supports later spatiotemporal reinstatement of prior experience.

To confirm the robustness of this finding beyond the sample of participants with epilepsy studied here, we analyzed the relation between revisitation and subsequent fixation-sequence replay in three independent datasets collected from neurologically typical individuals *(N* = 146 total, see Methods). Revisitation fixations during encoding significantly predicted later forward and reverse fixation-sequence replay (+1 and −1 lags) in each of the three datasets (Fig. 2c) (see Supplementary Table 1 for details). This establishes revisitation as a robust viewing pattern during encoding that supports the formation of spatiotemporal memory for scenes.

**Fig. 2.**
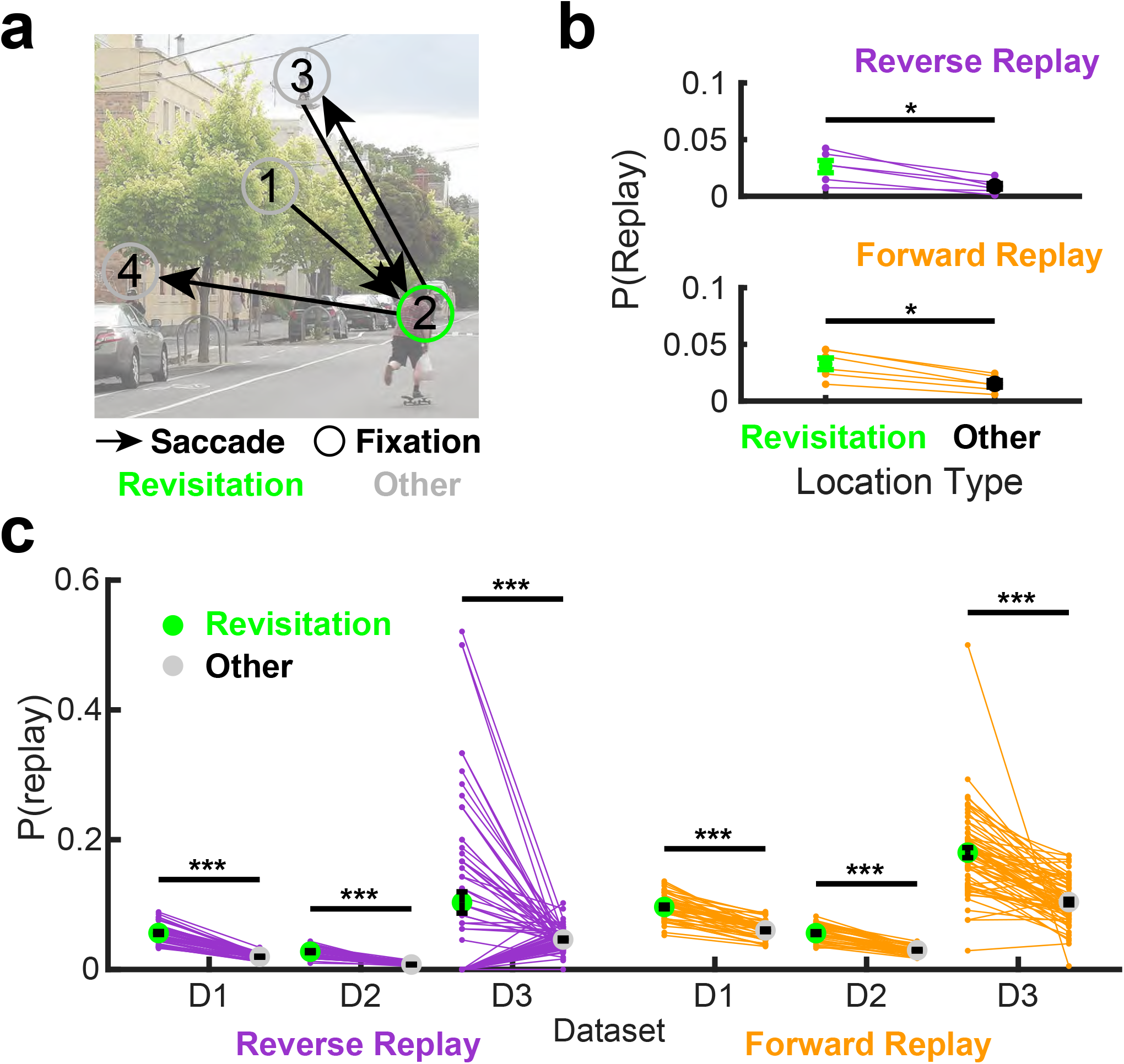
Revisitation enhances spatiotemporal memory formation. **a**. Revisitation fixations occurred during encoding when participants looked back at the previously viewed location as opposed to moving on to a new location. **b**. Relative to other fixations at encoding, participants were more likely to replay fixation sequences to revisited locations during retrieval, indicating that revisitation enhanced spatiotemporal memory formation. **c**. This relationship between revisitation at encoding and fixation-sequence replay at retrieval was robust in three independent datasets (D1-D3) collected from neurologically typical adults. Error bars denote SEM. *, *P* < 0.05. ***, *P* < 0.001. Dots indicate individual participants.

We next addressed the key question of whether activity of the hippocampus predicts the onset of revisitation fixations, as would be expected if it contributes to their generation. We focused on theta oscillations recorded from hippocampal iEEG depth electrodes (Fig. 3a), as theta is prominent in the field potential during memory-guided behaviors (3) and coordinates memory processing (25, 26). Hippocampal theta oscillations were prominent in field potentials recorded during task performance (Fig. 3b), with both low (< 4 Hz) and high (> 4 Hz) frequency theta oscillation peaks (Fig. 3c), often present at the same recording sites (Supplementary Fig. 2).

**Fig. 3.**
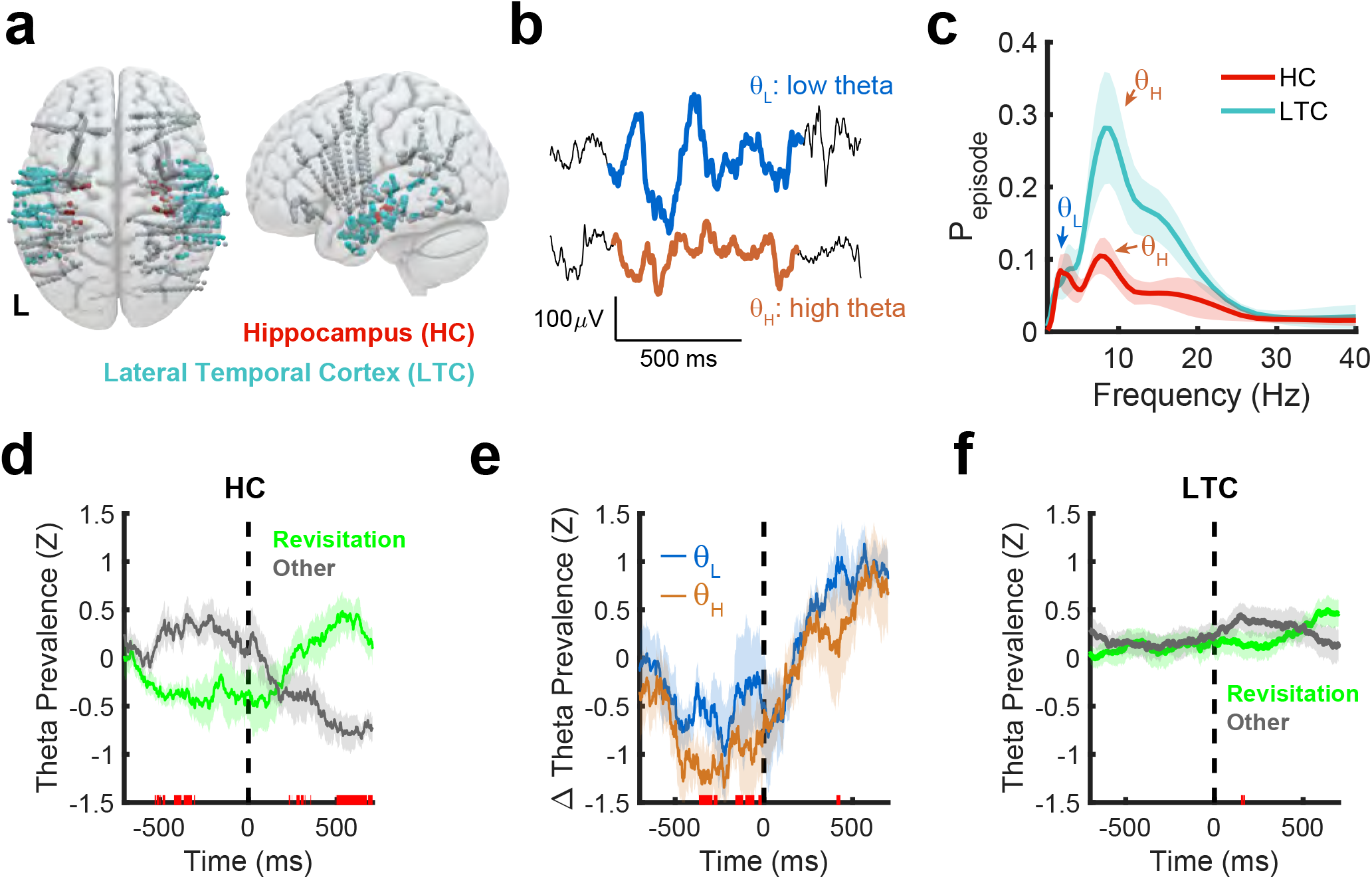
Revisitation is predicted by hippocampal theta oscillations. **a**. Field potentials were recorded from iEEG electrodes in bilateral hippocampus (HC, red) and in a control region (lateral temporal cortex, LTC, cyan). **b**. Example low and high frequency theta oscillations recording from the hippocampus. **c**. Oscillation detection revealed prevalent low and high frequency theta oscillations within the HC and LTC. **d**. In HC, theta prevalence differed significantly (*P* < 0.05, FDR corrected) preceding (~500 to 300 ms) and following (~500 to 700 ms) revisitation fixations (time = 0 indicates fixation onset). **e**. This relationship was more robust for higher than lower frequency theta oscillations. **f**. Within LTC, no significant differences in theta prevalence were detected leading up to revisitation fixations. Shaded regions denote 1 SEM across *N* = 6 participants.

Modulation of hippocampal theta oscillations prior to revisitation fixations would support an active role of hippocampus in their generation. In contrast, if the hippocampus were merely to respond to the perceptual input provided by revisitation fixations, then modulation of theta oscillations would be expected only in the post-fixation interval. We found evidence for hippocampal theta modulation related to revisitation in both the pre- and post-fixation intervals (Fig. 3d). From 528 to 300 ms before fixation onset, the prevalence of theta oscillations was significantly less for revisitation than for other fixations (*t*(5) = —3.5, *p* = 0.02, *g* = —2.0, 95% CI, —3.9—0.5). The opposite pattern followed fixation onset, with significantly greater theta prevalence from 495 to 700 ms for revisitation than other fixations (*t*(5) = 4.6, *p* = 0.006, *g* = 2.3, 95% CI, 0.9–4.2). Differences between revisitation and other fixations in the pre-fixation interval were significantly more pronounced for high than low theta (Fig. 3e, *t*(4) = 4.6, *p* = 0.01, *g* = 0.7, 95% CI, 0.2–1.5), whereas high and low theta were similarly affected by revisitation in the post-fixation interval. A control analysis indicated that pre-fixation differences in theta for revisitation versus other fixations were not driven by modulation of theta by the preceding fixation type (Supplementary Table 2).

Lateral temporal cortex (LTC) served as a control site, as theta oscillations in this region behave similarly to hippocampal theta (27), but we did not hypothesize that it would contribute to revisitation. Theta oscillations were robust in LTC, as expected (Fig. 3c). Theta oscillations in the LTC did not predict revisitation fixations (Fig. 3f) demonstrating a degree of anatomical specificity for the evidence of hippocampal theta predicting revisitation fixations.

We hypothesized that if hippocampal activity drives revisitation fixations, then the hippocampus should orchestrate activity within cortical regions that guide eye movements and process the information sampled by fixations. We expected that hippocampal activity prior to other (non-revisitation) fixations would reflect bottom-up influences from such regions, reflecting attentional control to sample visual information that feeds forward into hippocampus (28, 29). In contrast, to the extent that revisitation fixations were driven by hippocampal-dependent memory, we expected a shift towards top-down influences from the hippocampus to such regions. We focused our analysis to electrodes within two visually oriented networks (Fig. 4a): the dorsal attention network (DAN) which supports spatial attention and oculomotor control (29) and a visual network (VN) involved in perception. DAN and VN electrodes exhibited theta oscillations, with peak frequencies at 8.8 and 8.2 Hz, respectively (Fig. 4b). These theta oscillations synchronized with hippocampal theta during encoding, for both the VN (*t*(4) = 3.2, *p* = 0.03, *g* = 1.4 (95% CI, 0.1–2.7) and DAN (*t*(4) = 8.4, *p* = 0.001, *g* = 3.7 (95% CI, 1.1–6.4) (Fig. 4c,d). These findings parallel demonstrations of visual theta in rodents (30) and nonhuman primates (31) and suggest the hippocampus is linked to rhythmic perception and attention (32, 33).

**Fig. 4.**
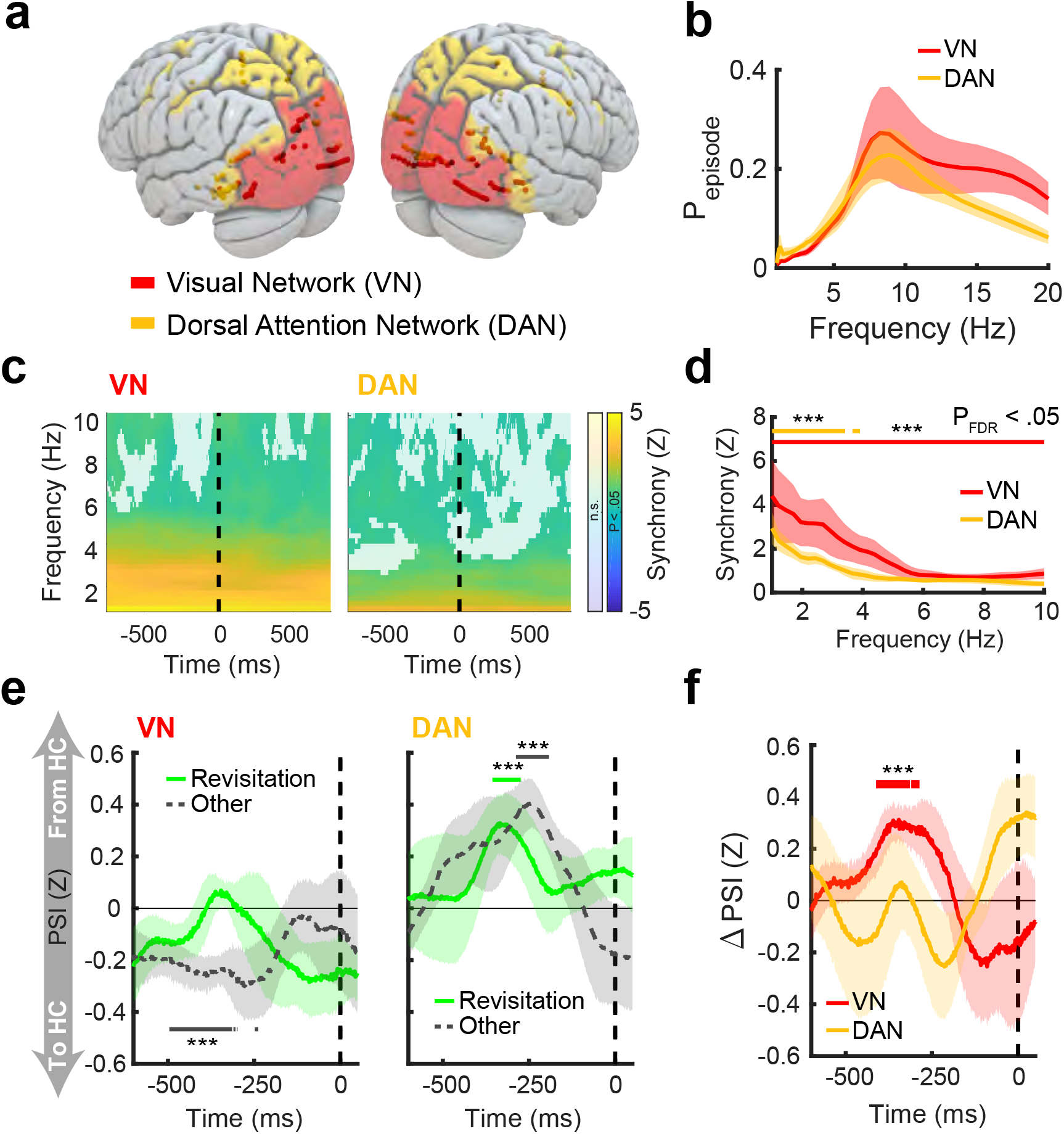
Directed influence of hippocampus on cortical network theta oscillations before fixations. **a**. Electrode locations in the DAN and VN regions of interest. **b**. Both networks exhibited peak oscillations (*P_episode_*) in the high theta range. **c**. Time-frequency plots indicate theta synchrony of hippocampus with the DAN and VN during encodin (time = 0 indicates fixation onset). A transparency mask highlights significant time-frequency points (*P* < .05, FDR corrected). **d**. Synchrony was consistently high across the pre- and post-fixation interval. **e**. Directional theta interactions between the hippocampus and cortical networks indicated that information flow (PSI) was directed from the VN to the hippocampus preceding non-revisitation fixations (left). Information flowed from the hippocampus to the DAN preceding both revisitation and other fixations (right). **f**. Information flow from the VN to the hippocampus was significantly greater preceding revisitation fixations than other fixations. All plots depict data from *N* = 5 participants with electrode contacts in these cortical networks and the hippocampus. Shaded regions depict standard error of the mean. ***, *P* < .05, FDR corrected.

We next tested the hypothesis that the hippocampus drives activity of these systems and that it does so to a greater extent during revisitation than other fixations. We calculated the phase slope index (PSI), a measure of directed information flow, between hippocampus and both systems. For the VN, there was significant bottom-up information flow to the hippocampus before participants made other (non-revisitation) fixations (significantly negative PSI (*t*(4) = −4.6, *p* = 0.01, *g* = −2.1 (95% CI, −3.7–-0.4) (Fig. 4e) that shifted to more top-down information flow from hippocampus to visual system before revisitation fixations (Fig. 4g, *t*(4) = 4.7, *p* = 0.009, *g* = 2.0 (95% CI, 0.6–3.9. We observed significant information flow from hippocampus to DAN before both revisitation fixations (*t*(4) = 2.9, *p* = 0.04, *g* = 1.3 (95% CI, 0.03–2.5) and other fixations (*t*(4) = 4.5, *p* = 0.01, *g* = 2.0 (95% CI, 0.4–3.6) fixations, without modulation by fixation type (Fig. 4f). Thus, hippocampal coordination of DAN occurred before all fixations whereas reversal of the typical bottom-up flow of information from the visual system to the hippocampus uniquely occurred before revisitation fixations.

Previous findings of the hippocampal involvement in guiding eye movements by long-term memory (7–9, 34) support the standard model of hippocampal long-term memory function (1). In contrast, the current focus on initial learning permitted testing of a far more immediate hippocampal contribution to short-term (i.e., within-episode) memory. We identified hippocampal contributions to memory processing across rapid gaze fixations, highlighting an active, immediate role of the hippocampus to guide forthcoming fixations in a manner conducive to learning. Revisitation fixations provided a temporally precise behavioral marker of this process. Revisitation fixations countered the typical pattern of looking towards novel rather than previously viewed scene content (24), suggesting sporadic guidance by memory retrieval, and enhanced learning, as indicated by better subsequent spatiotemporal memory. This interpretation is consonant with shifts in hippocampal states from pronounced theta oscillations during novel exploration, to theta-free epochs marked with sharp-wave ripple events (35) thought to support replay of prior experience (36). Eye-movement tracking therefore provided a marker of memory processing with the requisite temporal precision to resolve the behavioral ramifications of dynamic changes in hippocampal activity reflecting encoding versus retrieval that occur rapidly over a brief interval (34, 37), even during a single episode during which learning occurs for a novel stimulus.

These findings also situate the hippocampus within a distributed system for visual cognition, attention, and memory. The expected bottom-up flow of information from the visual stream to hippocampus was abated immediately before revisitation fixations. Furthermore, the cortical oculomotor system that guides visuospatial attention (29) is thought to be driven by perceptual and semantic scene features (28, 38). We identified a directed influence of hippocampal theta activity on this network preceding fixations of all types. These findings demonstrate hippocampal top-down influences on networks that serve perception and visuospatial attention, as previously hypothesized (11, 39). Because these hippocampal influences occurred during initial scene viewing, they cannot be explained by long-term memory, but rather must reflect the active guidance of attention and perception by short-term memory. The long and the short of hippocampal function may therefore be its critical role in the effective coordination of attention and perception during learning, explaining why hippocampal damage and dysfunction impairs long-term memory (1) as well as perception and short-term retention (15–18).

## ACKNOWLEDGEMENTS

Wethankthe patients of the Northwestern University Comprehensive Epilepsy Center and their families for their efforts to facilitate this research. This work was supported in part by National Institute of Neurological Disorders and Stroke grant T32NS047987.

## AUTHOR CONTRIBUTIONS

Conceptualization, J.E.K. and J.L.V., Formal analysis, J.E.K., Data Collection, J.E.K. and J.L.V., Writing – original draft preparation, J.E.K. Writing – review and editing, J.E.K., S.V.H., S.S., J.M.R., and J.L.V., Resources – S.V.H., S.S., J.M.R. and J.L.V., Supervision, J.L.V.

## Methods

### Participants

We enrolled six participants (3 female) with medically refractory epilepsy from the Northwestern Memorial Hospital Comprehensive Epilepsy Center (Chicago, IL). All participants had depth electrodes implanted as part of neurosurgical monitoring prior to elective surgery. Inclusion criteria for this study were implantation of electrodes into the hippocampus. The average age of participants was 29 (range 24 – 38) years. Written informed consent was acquired prior to participation in the research protocol in accordance with the Northwestern University Institutional Review Board.

### iEEG recordings

Stereotactic EEG electrodes (contacts spaced 5-10 mm apart, AD-TECH Medical Instrument Co., Racine, WI) targeted brain structures based on clinical needs, but provided coverage in the hippocampus as well as the dorsal attention and visual networks beyond the seizure onset zone. Electrophysiological data were recorded with a clinical reference and ground consisting of a surgically implanted electrode strip facing the scalp. Recordings were made using a Nihon Kohden amplifier with a sampling rate of 1–2 kHz, per clinical needs. Recorded signals were bandpass filtered from 0.6 to 600 Hz and re-referenced offline to a bipolar montage computed using adjacent electrode contacts. Line noise and harmonics (60, 120 and 180 Hz) were removed with a discrete Fourier transform filter. To rule out the possibility that epileptiform activity influenced our oscillation detection analyses, all data within 1 sec of inter-ictal epileptiform discharges were excluded from analysis using an automated algorithm that detects large amplitude increases in high frequency activity (40).

### Electrode localization

Post-implant CT were coregistered to presurgical T1 weighted structural MRIs using SPM12 (41). All T1-weighted MRI scans were normalized to MNI space by using a combination of affine and nonlinear registration steps, bias correction, and segmentation into grey matter, white matter, and cerebrospinal fluid components. Deformations from the normalization procedure were applied to individual electrode locations identified on post-implant CT images or structural images using Bioimage Suite (https://medicine.yale.edu/bioimaging/suite/). Bipolar pairs with at least one contact within either hippocampus, the dorsal attention network, or the visual network (Fig. 4) were analyzed. Electrode contacts in hippocampus were verified by visual inspection of coregistered anatomical scans and additionally included contacts directly adjacent to hippocampal grey matter in the hippocampal-amygdala transition area in 2 participants. Contacts in the DAN and VN were defined based on cortical parcellations of Yeo and colleagues (42). Maximum probability estimates from the Harvard-Oxford atlas (43) defined contacts within the lateral temporal cortex (see Supplementary Fig. 3 for additional information).

### Eye tracking

We recorded eye movements at 500 Hz using an Eyelink 1000 remote tracking system (SR Research, Ontario, Canada). Continuous eye-movement records were parsed into fixation, saccade, and blink events. Motion (0.15°), velocity (30°/*s*) and acceleration (8000°/*s*^2^) thresholds indicated saccade events. Blinks were determined by pupil size, and remaining epochs below detection thresholds were classified as fixations. The location of each fixation event was computed as the average gaze position throughout the duration of the fixation. The eye-tracking camera was mounted beneath the computer monitor that displayed the task, which was affixed to a movable arm to allow positioning directly in front of the patient’s eyes at a distance of approximately 60 cm.

### Recognition memory task

A scene recognition task was designed using Presentation^®^ software (Version 18.0, Neurobehavioral Systems, Inc., Berkeley, CA). The task consisted of 8 blocks, in which the participant studied a sequence of 24 images followed by a recognition test. The images contained common objects in naturally occurring contexts (44) restricted to scenes containing people, animals, or food (8 scenes of each category per block). During encoding, scenes were displayed for 3 seconds, followed by a randomly jittered inter-trial interval of 0.8 to 1.2 seconds (uniformly distributed). We presented a centrally located fixation cross for 0.5 seconds before each scene appeared to alert the participant to the upcoming trial. Scene images were displayed on the computer monitor and subtended approximately 24 by 24 degrees of visual angle.

During recognition, participants viewed 24 novel and 24 repeated scenes from the previous encoding block in pseudorandom order. Novel scenes were matched for content (they contained the same type of objects) to the repeated scenes but were not previously viewed in the task. The recognition phase of the task used a gaze-contingent design to allow memory to guide visual sampling. Scene content was masked by a grey overlay which was made transparent through a Gaussian kernel (*μ* = 0, *σ* =1.25 °of visual angle) centered at the current gaze position. An exponential moving average (*α* = 0.6) reduced transient shifts in the position of the revealed window. Prior to each recognition trial, a fixation cross appeared for 0.5 seconds at a cued location that indicated where to begin visual search. For a given scene, the cue appeared at the center of an object with either the highest or lowest visual salience, as defined by the DeepGaze II model (45). Gaze-contingent search followed until the participant indicated by button press that the scene was either repeated or novel. Recognition trials were separated by a 0.8 to 1.2 second inter-trial interval. Because behavioral performance (*d*’) did not differ between high and low salience cues (*t*(5) = −1.0, *p* = 0.36, *g* = −0.1 (95% CI, −0.4–0.2), we report analyses that collapse across these two conditions.

Throughout the task, eye-tracking validation was performed before encoding and recognition phases of every block. If the average error exceeded 1.5° of visual angle on the five-point calibration test, the eye tracker was re-calibrated before continuing with the task.

### Eye movement analysis

To measure replay of fixation sequences during scene recognition, we computed lag conditional viewing probability (lag-CVP) curves, a recently developed measure of gaze reinstatement (21). Briefly, gaze fixations made during encoding was serialized. Reinstatement during retrieval was measured by computing the probability of viewing a region of the scene, conditional on the distance (lag) of the previous fixation in the encoding sequence. Fixations at retrieval were matched to the encoding fixations using a nearest neighbor approach (using Euclidean distance), with a threshold of 2° of visual angle. Lags were computed based on this updated sequence. Replay was defined as lag 1 transitions in forward (+1) and reverse (—1) directions.

Revisitation fixations were defined as gaze fixations made during scene encoding when the gaze returned to a previously fixated location. As in the definition of reinstatement, a threshold of 2° of visual angle was used to assign fixations to specific locations. To examine the influence of revisitation on replay, fixated locations at study were labeled as involved in subsequent forward (+1 lags) or reverse (—1 lags) replay, or not. Then, the probability of replay was compared for revisited and non-revisited locations. To generalize our behavioral findings outside the sample of epileptic patients, we replicated these analyses in three independent datasets that used concurrent eye tracking with repeated viewing of scenes (46–48).

### Analysis of theta oscillations

As observed in humans, nonhuman primates, and bats (49–51), theta oscillations were transient in nature (Fig. 3b). We therefore quantified theta oscillations using *Pepisode,* which indicates the probability of an oscillation being present, using an automated oscillation detection algorithm (52, 53). Following this approach, we defined oscillatory episodes as temporal epochs that exhibited high spectral power within a specified frequency band that was sustained for at least three full cycles. Estimates of spectral power were obtained through Morlet wavelet convolution (6 cycles) at 50 frequencies from 1 to 40 Hz. We fitted a linear regression between the log of power and frequency across the entire recording session. The mean power for a given frequency was used to build a *χ*^2^ (2) distribution to represent the contribution of 1 /*f* signal to power at a given frequency. Epochs with spectral power above 95% of this distribution and duration of at least three cycles were considered oscillations. As theta oscillations were prevalent at lower (2-4 Hz) and higher (6-8 Hz) frequencies (Fig. 2b), we selected the peak theta frequency in each band, if present at a given electrode contact. Having identified bouts of theta oscillations, we measured the proportion of time that oscillations were present (*Pepisode*) for different event types of interest (53). *Pepisode* was then used as a dependent measure for statistical analysis.

### Directed coupling between hippocampus and visual systems

We measured synchrony between LFPs recorded from the hippocampus and visual regions using the phaselocking value (PLV) (54). For each pairwise combination of contacts in the hippocampus and cortical regions, we computed synchrony as described above. We standardize these measures across electrode contacts for group comparison using a permutation approach *(N* = 1000). We constructed a null distribution by shuffling the hippocampal phase estimates across trials and z-transformed observed values based on the mean and standard deviation of this null distribution. Prior to group analysis, all connections between the hippocampus and a given cortical region of interest were averaged. Statistical analysis was then performed using nonparametric tests as described below.

After establishing synchrony between the hippocampus and cortical sites, we tested for potential directional interactions using the phase slope index (PSI) (55). We computed the PSI in a time-resolved manner in the moments surrounding both revisitation and other fixations. For two recorded signals, PSI is based on the slope of phase differences (between two recorded sources) with increasing frequency. To standardize PSI values, phase values were permuted (*N* = 1000) and a distribution was generated to estimate the mean and standard deviation under the null hypothesis of no relation between frequency and phase differences. These standardized scores were then submitted to non-parametric permutation statistics, as described below.

### Quantification and statistical analysis

Data are reported as mean ± standard error. Non-parameteric (Wilcoxon signed rank, Wilcoxon rank sum, Kruskal Wallis) statistical tests were used for tests on bounded observations (e.g., prevalence of oscillations). Parametric tests (paired or one sample t tests) were applied to data with normally distributed residuals. Nonparametric permutations tests (56) with FDR correction for multiple comparisons (57) were applied for all timeseries and time-frequency analyses. Alpha was set to 0.05 for determining statistical significance. Repeated measures ANOVAs were used for behavior analysis. Hedges’ g (58) was used to estimate effect sizes, with values estimated at peaks for timeseries analysis. For such tests involving multiple comparisons, these estimates are necessarily biased and serve descriptive purposes only.

## Supplementary Tables

**Supplementary Table 1.** Effect of revisitation sampling on fixation-sequence replay.

|  |  | Revisitation | Other | t-value | df | g | p |
| --- | --- | --- | --- | --- | --- | --- | --- |
| Dataset 1 (47) | Forward Replay | 0.10 (0.02) | 0.06 (0.01) | 14.1 | 44 | 2.1 | <.0001 |
|  | Reverse Replay | 0.06 (0.01) | 0.02 (0.005) | 19.3 | 44 | 3.3 | <.0001 |
| Dataset 2 (48) | Forward Replay | 0.06 (0.01) | 0.03 (0.007) | 15.0 | 63 | 2.6 | <.0001 |
|  | Reverse Replay | 0.03 (0.01) | 0.01 (0.003) | 13.1 | 63 | 3.1 | <.0001 |
| Dataset 3 (46) | Forward Replay | 0.18 (0.06) | 0.10 (0.03) | 9.0 | 66 | 1.5 | <.0001 |
|  | Reverse Replay | 0.10 (0.14) | 0.05 (0.02) | 3.5 | 65 | 0.6 | .0009 |

The probability of replay was compared for revisitation and other fixations. Revisitation fixations led to increased fixation-sequence replay across all three datasets.

**Supplementary Table 2.** Control analyses predicting revisitation fixations from theta oscillations.

|  |  | t-value | df | Hedges' g (95% CI) | p |
| --- | --- | --- | --- | --- | --- |
| Revisitation Control | Pre-fixation | -4.1 | 5 | -2.4 (-4.5 – -0.8) | .009 |
|  | Post-fixation | 5.5 | 5 | 2.7 (1.2 – 4.8) | .003 |
| Other Control | Pre-fixation | -3.6 | 5 | -2.0 (-3.0 – -0.6) | .01 |
|  | Post-fixation | 5.6 | 5 | 2.4 (1.1 – 4.3) | .003 |

Reported statistic tests control for sequential viewing behaviors, by excluding fixations that are preceded by either revisitation or non-revisitation fixations, which are accompanied by relative decreases and increases in theta oscillations in the pre-fixation interval. The reported summary statistics are averaged across all timepoints in the pre- or post-fixation interval that survived multiple comparison correction (*P* < .05, FDR corrected).

## Supplementary Figures

**Supplementary Figure 1.**
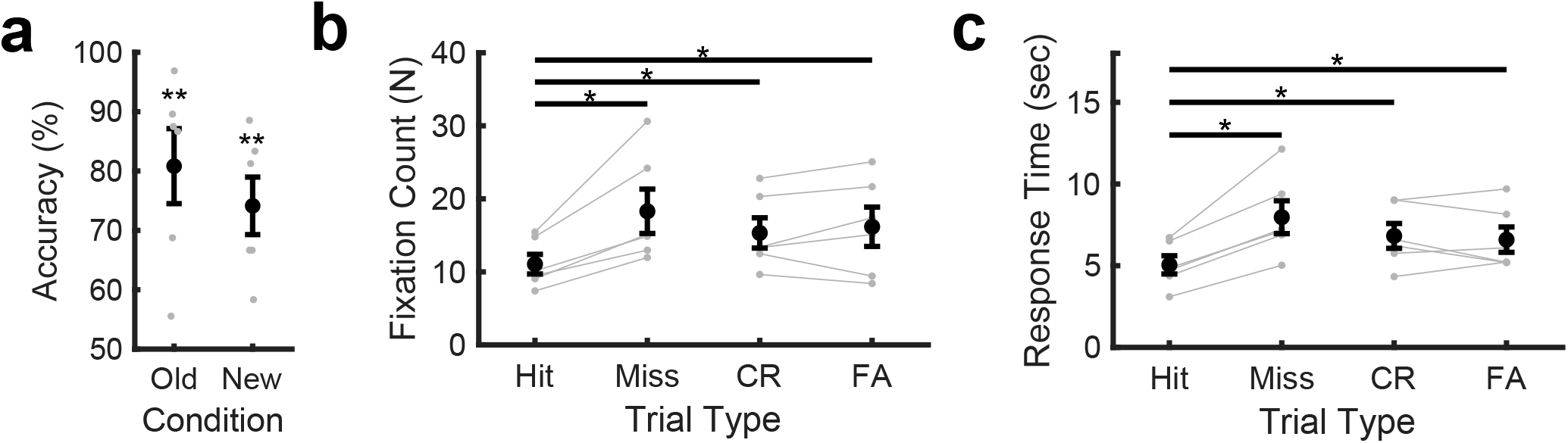
Recognition accuracy and relationship between recognition and eye-movement behavior. **a**. Participants accurately recognized previously viewed (old) and completely novel (new) scenes during the test phases. **b**. Accurate recognition of old scenes (hits) involved significantly fewer gaze fixations than recognition misses, correct rejections (CR) of new items, and false alarms (FA) to old items. **c**. Recognition memory accuracy was related to recognition response times in the same manner, indicating memory-guided visual search determined responding. **, *P* < .001. *, *P* < .05. Error bars denote SEM. Dots indicate individual participants.

**Supplementary Figure 2.**
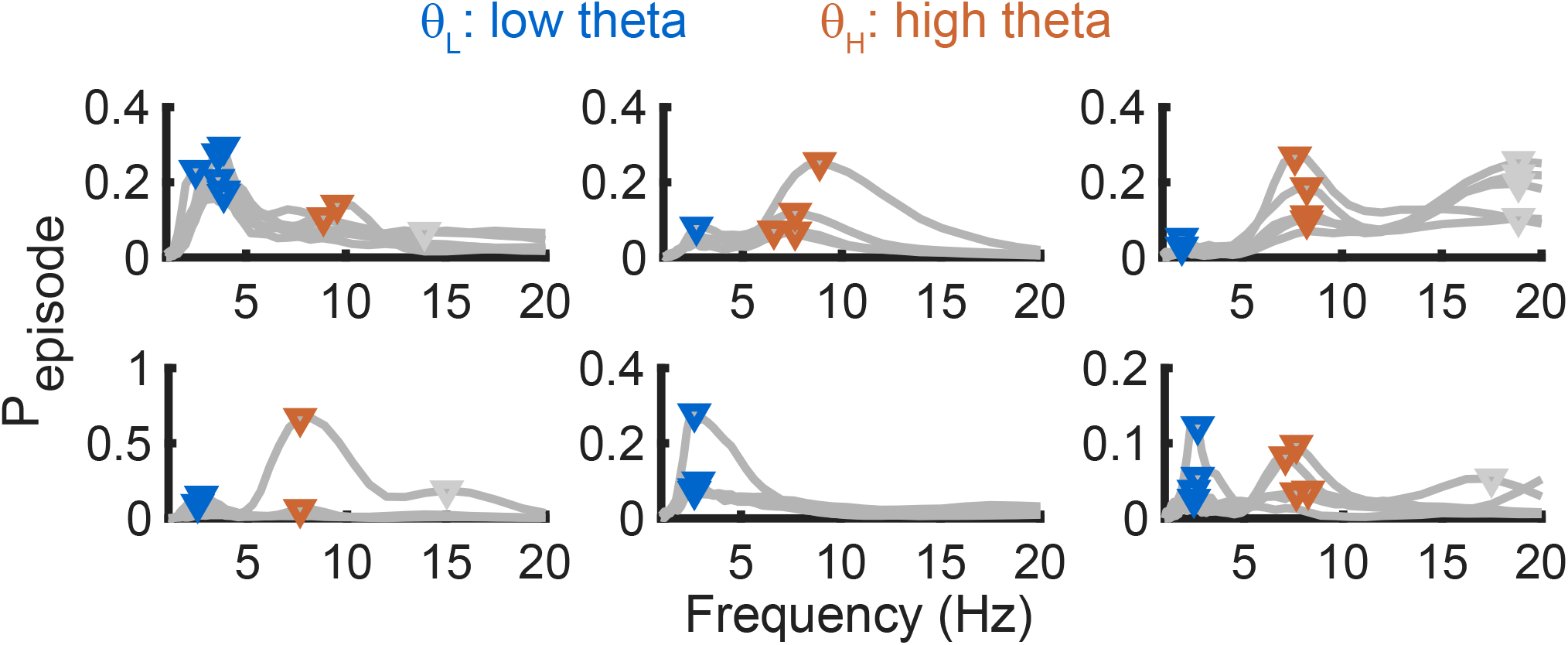
Prevalence of hippocampal theta oscillations for individual participants and electrodes. Low frequency (2 — 5 Hz) theta oscillations were detected in all 6 participants and high frequency (5 — 10 Hz) theta oscillations were detected in all but one participant. Curves denotes oscillation prevalence for individual electrodes, plotted separately for each participant.

**Supplementary Figure 3.**
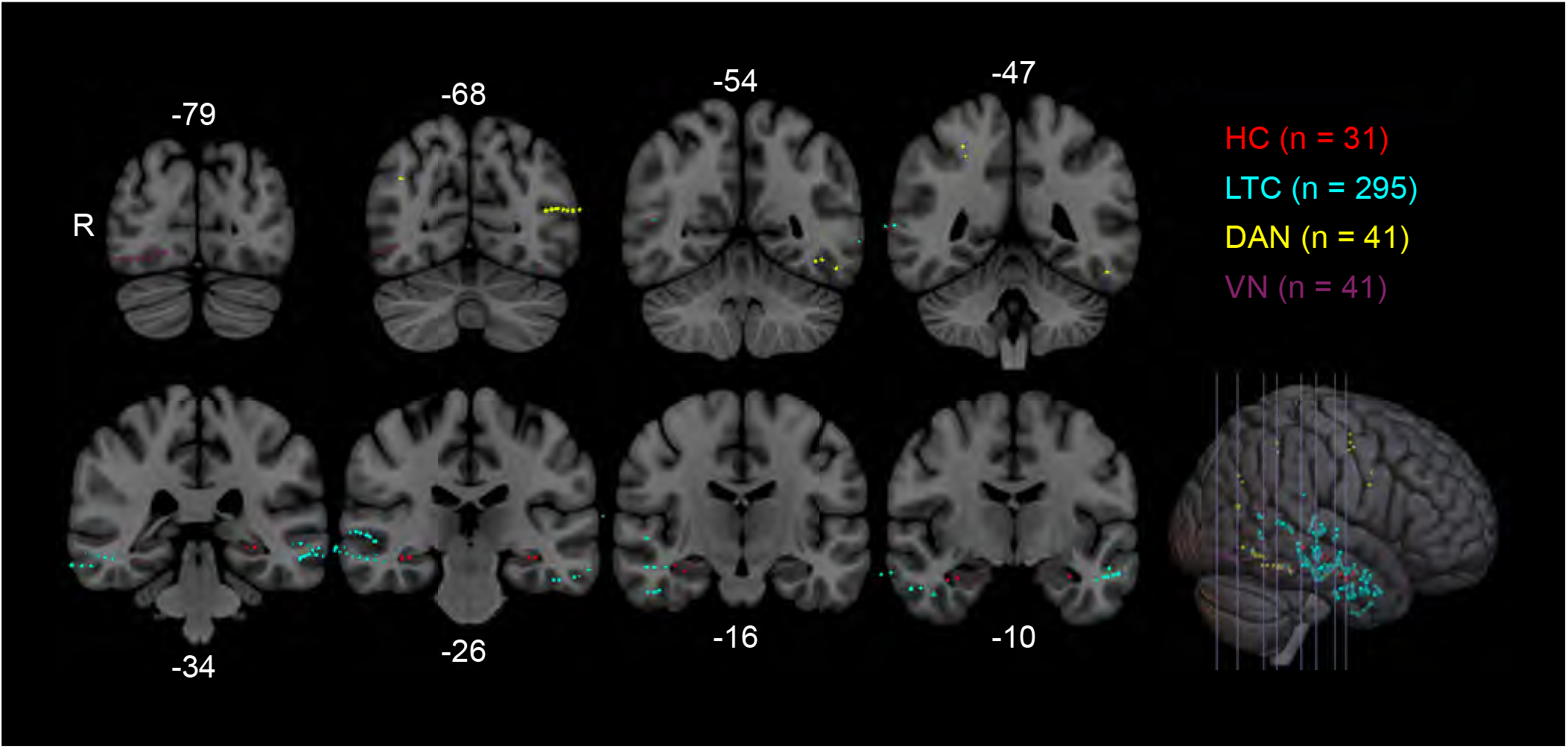
Electrode coverage in regions of interest. Locations of electrode contacts within the hippocampus (HC, 4 — 6 per participant), lateral temporal cortex (LTC, 37 — 57 per participant), dorsal attention network (DAN, 2 — 14 per participant) and visual network (VN, 1 — 13 per participant) are depicted on coronal slices in MNI space (labels denote the y coordinate, in mm) overlaid on the ICBM 2009b Nonlinear Asymmetric template (59). Slices are displayed in radiological convention (contacts in the right hemisphere are display on the left side of the brain.

